# *Nos2*^-/-^ mice infected with *M. tuberculosis* develop neurobehavioral changes and immunopathology mimicking human central nervous system tuberculosis

**DOI:** 10.1101/2021.08.15.456427

**Authors:** Xuan Ying Poh, Jia Mei Hong, Chen Bai, Qing Hao Miow, Pei Min Thong, Yu Wang, Ravisankar Rajarethinam, Cristine S.L. Ding, Catherine W.M. Ong

## Abstract

**Background:** Understanding the pathophysiology of central nervous system tuberculosis (CNS-TB) is hampered by the lack of a good pre-clinical model that mirrors the human CNS-TB infection. We developed a murine CNS-TB model that demonstrates neurobehavioral changes with similar immunopathology with human CNS-TB.

**Methods:** We injected two *Mycobacterium tuberculosis* (*M*.*tb*) strains, H37Rv and CDC1551, respectively, into two mouse strains, C3HeB/FeJ and *Nos2*^-/-^ mice, either into the third ventricle or intravenous. We compared the neurological symptoms, histopathological changes and levels of adhesion molecules, chemokines, and inflammatory cytokines in the brain induced by the infections through different routes in different strains.

**Results:** Intra-cerebroventricular infection of *Nos2*^-/-^ mice with *M*.*tb* led to development of neurological signs and more severe brain granulomas compared to C3HeB/FeJ mice. Compared with CDC1551 *M*.*tb*, H37Rv *M*.*tb* infection resulted in a higher neurobehavioral score and earlier mortality. Intra-cerebroventricular infection caused necrotic neutrophil-dominated pyogranulomas in the brain relative to intravenous infection which resulted in disseminated granulomas and mycobacteraemia. Histologically, intra-cerebroventricular infection of *Nos2*^*-/-*^ mice with *M*.*tb* resembled human CNS-TB brain biopsy specimens. H37Rv intra-cerebroventricular infected mice demonstrated higher brain concentrations of inflammatory cytokines, chemokines and adhesion molecule ICAM-1 than H37Rv intravenous-infected mice.

**Conclusions:** Intra-cerebroventricular infection of *Nos2*^-/-^ mice with H37Rv creates a murine CNS-TB model that resembled human CNS-TB immunopathology, exhibiting the worst neurobehavioral score with a high and early mortality reflecting disease severity and its associated neurological morbidity. Our murine CNS-TB model serves as a pre-clinical platform to dissect host-pathogen interactions and evaluate therapeutic agents for CNS-TB.

## BACKGROUND

The most severe form of *Mycobacterium tuberculosis* (*M*.*tb*) infection is central nervous system tuberculosis (CNS-TB) which has high mortality and serious long-term neurological sequelae even with effective anti-tuberculous treatment [1-3]. Common manifestations of human CNS-TB are tuberculous meningitis (TBM), tuberculomas and tuberculous brain abscesses [4]. Cerebral vasculitis and inflammation resulting in infarcts is the primary cause of permanent brain tissue damage in TBM and is among the worst consequences of CNS-TB [5, 6]. Despite effective TB treatment with antibiotics and adjunctive corticosteroids, CNS-TB remains one of the more challenging clinical syndromes to manage.

To advance our understanding of CNS-TB, we need an appropriate animal model that recapitulates the neurobehavioral, immunopathological and histopathological changes in human CNS-TB to dissect pathogenesis and aid drug discovery. Several animal models of CNS-TB have been described, including guinea pigs, rabbits, mice, pigs, and zebrafish. The rabbit model closely mimics human disease, developing clinical and histological changes [7-13]. However, a number of immunological tools profiling protein secretion and gene expression are unavailable for rabbits [1] and therefore preclude their suitability for in-depth immunological studies.

The mouse model has many advantages over other animals, including the availability of genetic and molecular tools as well as cost-effectiveness for large studies. However, existing murine CNS-TB models do not display the clinical features and immunological phenotypes of CNS-TB observed in humans. C57BL/6 mice are generally resistant to CNS-

TB infection, with no pathological abnormalities detected and no observed mortality over 24 weeks of study [14]. BALB/c mice infected through the intracerebral route directly into the brain parenchyma with *Mycobacterium bovis* BCG (BCG) had infiltration of inflammatory cells, but no granulomas were observed [10]. This contrasts with human CNS-TB where tuberculomas occur in approximately 30% of TBM patients [15]. Intravenous inoculation of BALB/c mice with *M*.*tb* strain CDC1551 successfully infected the CNS but did not produce granulomas in the brain and had low expression of brain chemokines and cytokines IL-1β, IL-6, TNF-α and IFN-γ, in contrast to the increased expression of these cytokines in the cerebrospinal fluid (CSF) of human TBM patients [16, 17]. While some murine CNS-TB models have meningitis and/or brain granulomas, they do not demonstrate neurological signs of disease and mortality, unlike human CNS-TB [14, 18]. Given the varying susceptibility and pathology of CNS-TB infection in different mouse strains, genetic predisposition is likely to play a crucial role. C3HeB/FeJ “Kramnik” mice were found to be hyper susceptible to *M*.*tb* infection due to the presence of an allele, termed the “super susceptibility to tuberculosis 1” (*sst1*) locus, and developed a more human-like lung pathology in contrast to C57BL/6 mice [19, 20]. However, the ability of C3HeB/FeJ mice to develop CNS-TB remains to be explored.

Intracerebral-infection with *M*.*tb* H37Rv directly into the brain parenchyma of inducible nitric oxide synthase (iNOS)-knockout mice resulted in neurological signs with meningitis, and mice exhibited 63% mortality post-infection (p.i.) [21]. However, the development of intracerebral tuberculomas and immunological profile were not phenotyped in this mouse model. Cytokine-induced upregulation of iNOS or NOS2 by murine macrophages have been implicated in the killing of intracellular pathogens such as *M*.*tb*, but the role of this antimicrobial system in human macrophages remains unclear [22, 23]. Studies have shown that activated human microglia, the brain resident macrophages, do not express iNOS [24, 25] or reactive nitrogen intermediate (RNI) nitric oxide (NO) [26], whereas murine microglia produced substantial amounts of NO on activation [26]. Given the well-established role of macrophages in TB, the inter-species difference in microglia expression of iNOS may explain the species tropism barrier to the development of CNS-TB in mice.

To address the limitations of existing murine CNS-TB models, we explored the effects of mouse strains, *M*.*tb* strains and routes of infection on the development of CNS-TB disease. First, we compared two mouse strains, C3HeB/FeJ and *Nos2*^-/-^ mice, to investigate whether the *sst1* locus or *Nos2* gene plays a more important role in CNS-TB infection. After selecting the suitable mouse strain, we investigated the effects of two different *M*.*tb* strains, H37Rv and CDC1551, and two routes of infection: intra-cerebroventricular (i.c.vent.) into the third ventricle and intravenous (i.v.), on the development of a murine CNS-TB model with human-like pathology. The i.c.vent. route of infection mimics the rupture of meningeal tuberculous lesions and the subsequent release of *M*.*tb* into the CSF, whereas the i.v. route mimics the hematogenous spread of *M*.*tb*. In this study, we showed that i.c.vent. infection of *Nos2*^-/-^ mice with *M*.*tb* H37Rv developed the severe neurological symptoms and induced a high expression of adhesion molecules, chemokines, and inflammatory cytokines in the brain, consistent with the infiltration of inflammatory cells and pathological changes. This pre-clinical model can be used to understand the mechanisms in host immunopathology and evaluate treatment for CNS-TB.

## METHODS

### Human CNS-TB brain specimen processing

The paraffin blocks of three surgical samples from patients with histological features indicative of CNS-TB infection were retrieved from the files of the Department of Pathology at Tan Tock Seng Hospital, Singapore. The specimens included leptomeninges and brain parenchyma, and demonstrated granulomatous inflammation typical of CNS-TB. Acid-fast bacilli were demonstrated on Ziehl-Neelsen histochemical stain in two out of three samples. Control brain sections were from the non-neoplastic brain parenchyma of three surgical pathology brain resection specimens. 4 μm thick sections were cut from each block for H&E staining according to the manufacturer’s instructions.

### Bacterial strains and growth conditions for infection

*Mycobacterium tuberculosis* (*M*.*tb*) strains H37Rv and CDC1551 were kindly provided by Professor Nick Paton and Associate Professor Sylvie Alonso (both NUS, Singapore) respectively. For infection experiments, a frozen vial of *M*.*tb* was thawed and cultured to mid-logarithmic phase at an optical density of 0.6-0.8. Prior to infection, the *M*.*tb* was centrifuged at 3,200 x g for 10 minutes and resuspended in 1 mL sterile 0.9% NaCl. The *M*.*tb* inoculum was then plated to determine the dose of infection.

### Mouse cannula implantation and infection

Six- to eight-week-old male C57BL/6 *Nos2*^-/-^ (Stock No. 002609) and C3HeB/FeJ (Stock No. 000658) mice (Jackson Laboratory, Bar Harbor, Maine) were used for intra-cerebroventricular (i.c.vent.) or intravenous (i.v.) infection. Mice in the i.c.vent. group were cannulated one week before infection. An illustration of the stereotaxic coordinates of site of injection and the positioning of guide cannula is shown in Supplementary Figure 1a. A motorized stereotaxic instrument (Neurostar, Tübingen, Germany) was used to implant a 26-gauge guide cannula (RWD, Shenzhen, China) into the third ventricle (coordinates from the bregma: -1.6 mm posterior, 0 mm lateral, -2.5 mm ventral). The same coordinates were used for both C57BL/6 *Nos2*^-/-^ and C3HeB/FeJ Kramnik mice as the size of mice were similar at the time of cannulation. *Nos2*^-/-^ mice were 23.5 g (± 1.1) (mean ± s.d.) and C3HeB/FeJ Kramnik mice were 24.7g (±1.6) (p = NS). Mice were injected with 0.5 μL of sterile 0.9% NaCl or 2 × 10^8^ CFU/mL *M*.*tb* through the brain cannula (over 5 min) using the syringe pump (Harvard Apparatus, Holliston, Massachusetts). Mice in the i.v. group were injected with 50 μL of sterile 0.9% NaCl or 2 × 10^6^ CFU/mL *M*.*tb* via the retro-orbital sinus. *M*.*tb* was administered at a dose of 10^5^ CFU to each animal, irrespective of the route of infection. This dose was chosen as previous CNS-TB murine models have administered *M*.*tb* within the range of 10^5^ to 10^6^ CFU [14, 16, 21, 27]. However, different infection routes have different recommended administration volumes (0.5 μL for i.c.vent. and 50 μL for i.v.) and the concentration of *M*.*tb* for i.c.vent. route was 100-fold more concentrated than the i.v. route. All mice were observed for mortality and weight change. Humane endpoints included ≥ 20% weight loss, complete hind limb paralysis and repeated seizures. Infected mice were also monitored daily for 56 days after infection for clinical signs indicative of CNS-TB, such as limb weakness, tremors, and twitches.

30 μL of trypan blue was administered into four cannulated *Nos2*^-/-^ mice and the brains harvested 15 mins post-administration to allow for distribution of the dye in both right and left cerebral hemispheres. A sagittal illustration of the ventricular system in the mouse brain, which include the lateral ventricles, third ventricle and aqueduct that leads to the fourth ventricle, is depicted in Supplementary Figure 1b. Coronal sections of each brain verifies that the dye is in the ventricular system (Supplementary figure 1c), indicating successful brain cannulation into the third ventricle. Given the similar sizes of both strains of mice at cannulation, trypan blue was not instilled into the ventricles of the C3HeB/FeJ Kramnik mice, but H&E staining of i.c.vent.-infected Kramnik mice showed more marked meningeal inflammation than the brain parenchyma (Supplementary Figure 1e), indicating the accurate placement of the cannula into the cerebral ventricles. *Nos2*^-/-^ or C3HeB/FeJ mice were infected with *M*.*tb* 7 days after brain cannulation, and the blood, brain, lungs, liver and spleen were harvested 56 days post-infection (p.i.) for enumeration of mycobacterial load, histopathological analysis and immunological marker analysis (Supplementary figure 1d).

### Neurobehavioral scoring

Neurobehavioral scoring was performed by a researcher (P.X.Y.) blinded to the route of infection and strain of *M*.*tb* according to a scoring list for CNS-TB mouse model (Table 2), adapted from Tucker et al [12]. Each scoring parameter ranged from one, corresponding to no abnormalities, to a variable maximum score. The minimum total score is 3 indicating a normal mouse. Higher neurological scores reflect an increasing severity of neurological deficits with a maximum total score of 7.

### Organ harvesting and processing

Eight weeks post-infection, mice were deeply anesthetized before cardiac puncture was performed for blood collection. The brain, lungs, liver and spleen were harvested and the gross pathology examined before tissue processing. Half of each organ was fixed in 10% neutral buffered formalin for histology, while the other half was homogenized for bacterial enumeration and characterization of immunological markers. Organs were homogenized by high-speed shaking in 2 mL microcentrifuge tubes filled with sterile PBS and 5/7 mm stainless steel beads using TissueLyser LT (Qiagen, Hilden, Germany).

### Histopathological analysis

Histopathology was performed on the left hemisphere of the murine brain. The murine brain was fixed in 10% neutral buffered formalin, paraffin embedded and sectioned to 4 μm thickness. Brain slices were stained with hematoxylin-eosin (H&E) (Vector Laboratories, Burlingame, California) to characterise pathological lesions and Ziehl-Neelsen staining (Sigma-Aldrich, St. Louis, Missouri) to detect mycobacterium according to manufacturer’s instructions. Histopathological examination was carried out in a blinded fashion by a histopathologist (R.R.) based on the presence of pathological changes including inflammation, perivascular cuffing, gliosis, neuronal necrosis, granuloma, pyogranuloma and necrosis. Definition of granulomatous lesions in this study includes both granulomas and pyogranulomas. Grading of severity was assigned on the following scale: 0: no abnormalities detected; 1-minimal; 2-mild; 3-moderate; 4-marked & 5-severe. The total number and area of granulomatous lesions were measured from 6 different sections of 5-6 mice. To quantify the area of granuloma, we utilized the free-hand tool in ImageJ (NIH, Bethesda, Maryland) and manually demarcated the granuloma as a region of interest for area measurement.

### Immunological marker analysis

Adhesion molecules, cytokines and chemokines were analysed by Fluorokine multianalyte profiling kit according to the manufacturer’s protocol (R&D Systems, Minneapolis, Minnesota) on the Bio-Plex 200 platform (Bio-Rad, Hercules, California). The minimum detection limit for the ICAM-1 and p-selectin were 52.7 pg/ml and 2.6 pg/ml respectively. The minimum detection limit for the cytokines and chemokines were CCL-2/MCP-1 134 pg/ml, CCL-3/MIP-1α 0.452 pg/ml, CCL-4/ MIP-1β 77.4 pg/ml, CCL-5/ RANTES 19.1 pg/ml, CCL-7/ MCP-3 1.69 pg/ml, CCL-8/ MCP-2 0.283 pg/ml, CCL-11/Eotaxin 1.46 pg/m, CCL-12/ MCP-5 0.613 pg/ml, CCL-19/ MIP-3β 2.28 pg/ml, CCL-20/ MIP-3α 3.95 pg/ml, CCL-22/ MDC 0.965 pg/ml, CXCL-1/ KC 32.9 pg/ml, CXCL-2/ MIP-2 1.97 pg/ml, CXCL-10/ IP-10 6.85 pg/ml, CXCL-13/ BLC 19.3 pg/ml, IL-1α 8.17 pg/ml, IL-1β 41.8 pg/ml, IL-6 2.30 pg/ml, IL-12 p70 12.8 pg/ml, IL-17A 7.08 pg/ml, IL-27 1.84 pg/ml, LIX 2.02 pg/ml, TNF-α 1.47 pg/ml, IFN-γ 1.85 pg/ml. Brain homogenates were assayed at neat for all analytes and results were normalised to their total protein concentrations (Bio-Rad, Hercules, California).

### Statistics

Continuous variables are presented as medians and interquartile range. Neurobehavior scores and body weight change between groups were compared using two-way ANOVA with post-hoc Tukey’s multiple comparisons test. Levels of adhesion molecules, cytokines and chemokines, and CFU counts between groups were compared using Mann-Whitney test. Comparison of survival curves between groups was calculated using the log-rank test. A two-sided value of *p* < 0.05 was considered significant. All analyses were performed using GraphPad Prism version 7.05 (Graphpad, San Diego, California).

## RESULTS

### *M*.*tb* infected *Nos2*^-/-^ strain exhibited worse neurobehavioral score and worse histopathological changes in the brain than C3HeB/FeJ strain

To investigate whether *Nos2*^-/-^ or C3HeB/FeJ mice better replicate human CNS-TB, we inoculated each mouse with 9.15 ± 2.33 × 10^4^ colony forming units (CFU; mean ± s.d) of *M*.*tb* CDC1551 into the third ventricle to infect the meninges (Supplementary figure 1). Infected *Nos2*^-/-^ mice displayed neurological symptoms such as twitching and limb weakness from 3 weeks post-infection (p.i.) (Video 2) that were not observed in infected C3HeB/FeJ mice or saline control mice (Video 1). Infected *Nos2*^-/-^ mice had significantly higher neurobehavioral scores than infected C3HeB/FeJ mice at 4 and 8 weeks p.i. (Figure 1a, p < 0.0001 and p < 0.0001 respectively). Neurological behavior assessed include tremors, twitches and appearance of eyes, with higher neurobehavioral scores reflecting an increasing severity of neurological deficits. CFU enumeration showed that brain and lung homogenates of infected *Nos2*^-/-^ mice had higher mycobacterial load compared to infected C3HeB/FeJ mice that had a trend to statistical significance (Figure 1b). Median (IQR) brain CFU count in *Nos2*^-/-^ and C3HeB/FeJ mice was 5 × 10^5^ (1.65 × 10^5^ – 5.8 × 10^5^) compared to 9.75 × 10^2^ (6.25 × 10^1^ – 5 × 10^3^) respectively (*p* = 0.057), while median (IQR) lung CFU count was 1.00 × 10^3^ (6.5 × 10^2^ – 1.5 × 10^3^) in infected *Nos2*^-/-^ mice and 0 (0 – 75) in infected C3HeB/FeJ mice (*p* = 0.057). Mycobacterial load in the liver, spleen and blood were similar.

**Figure 1.**
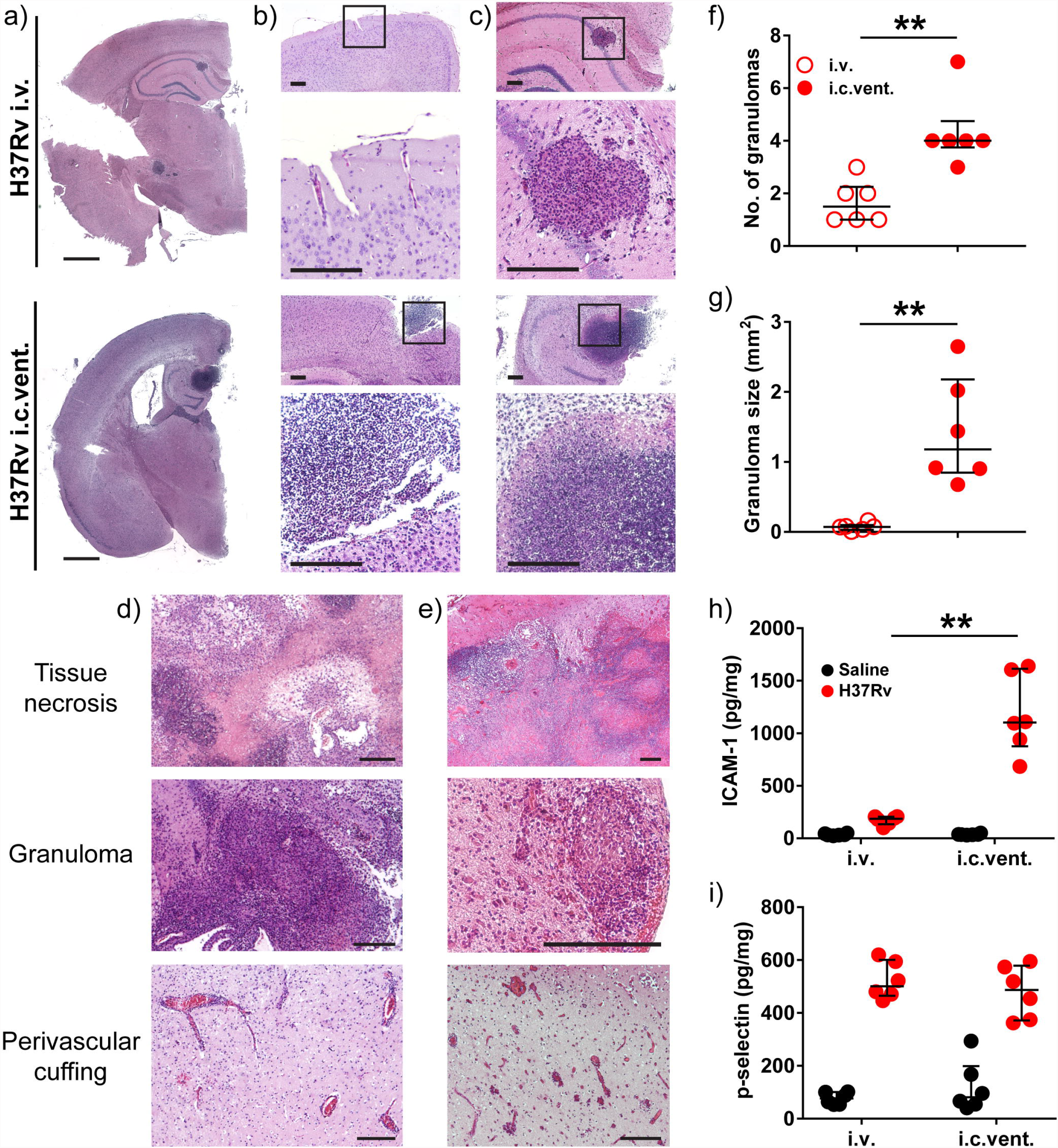
*Nos2*^-/-^ strain exhibited higher neurobehavioral score and increased inflammatory cell infiltrate in the brain compared to C3HeB/FeJ strain post-i.c.vent. infection with *M*.*tb* CDC1551. (a) Neurobehavioral scores were significantly higher in infected *Nos2*^-/-^ mice at 4 and 8 weeks p.i. compared with infected C3HeB/FeJ mice. Parameters assessed include tremors, twitches and appearance of eyes, with higher neurobehavioral scores reflecting an increasing severity of neurological deficits. Statistical analysis conducted using two-way ANOVA with post-hoc Tukey’s multiple comparisons test. ****, p < 0.0001. (b) *M*.*tb* colony forming units (CFU) in the brain and lung of *Nos2*^-/-^ is higher compared to C3HeB/FeJ mice at day 56 p.i., analyzed using Mann-Whitney test. (c) Macroscopic assessment of brain, lung and spleen of *M*.*tb* infected Nos2^-/-^ and C3HeB/FeJ mice were similar to saline controls. Images are representative of 2-4 mice per condition. Scale bar = 1 cm. (d) Hematoxylin and eosin (H&E) stain of a representative brain section from each group is shown, demonstrating normal brain histology in saline control mice and histopathology in infected mice. High-power views (insets) demonstrate more inflammatory cell infiltrate in the brain of infected *Nos2*^-/-^ compared to C3HeB/FeJ mice. Scale bar = 200 μm. (e and f) Infected *Nos2*^-/-^ have increased concentrations of (e) ICAM-1 and (f) p-selectin in the brain compared to C3HeB/FeJ mice. Adhesion molecule concentrations were normalised to total protein concentration and compared using two-way ANOVA with Sidak’s multiple comparisons test. Bars represent median and IQR. *, p < 0.05; **, p < 0.01; ***, p < 0.001. Inf = infected; Sal = saline. *Nos2*^-/-^ saline: n = 2; *Nos2*^-/-^ infected: n = 3; C3HeB/FeJ saline: n = 4; C3HeB/FeJ infected: n = 4.

Although there were no macroscopic changes in the brain, lung and spleen between the two mouse strains (Figure 1c), histopathological analysis revealed considerable differences between these two strains (Figure 1d). Infected *Nos2*^-/-^ mice demonstrated more inflammatory cell infiltrate in the brain parenchyma compared to infected C3HeB/FeJ mice. We postulated that the increase in leukocyte inflammation might be due to increased expression of adhesion molecules in the brain, and confirmed a significantly higher concentration of ICAM-1 and p-selectin in infected *Nos2*^-/-^ than C3HeB/FeJ mice (Figure 1e and f). Brain concentration of ICAM-1 and p-selectin were 14-fold (p = 0.0089) and 10-fold (p = 0.0008) higher in infected *Nos2*^-/-^ compared to C3HeB/FeJ mice.

Next, we investigated further the mechanism behind the increased immune cell recruitment in infected *Nos2*^-/-^ mice. As TB is characterised by a Th1 inflammatory response, we examined the concentrations of Th1 cytokines and chemokines. Concentrations of neutrophil chemoattractants were also profiled as histopathological analysis showed marked neutrophilic inflammation. Concentrations of Th1-associated inflammatory mediators TNF-α and CXCL-10 were significantly higher in infected *Nos2*^-/-^ mice than infected C3HeB/FeJ mice, while IFN-γ and CCL-5 showed a trend to increase (Figure 2a, b, c and d). Infected *Nos2*^-/-^ mice also had a significantly higher concentration of chemoattractants, CXCL-1, CXCL-2 and LIX, than infected C3HeB/FeJ mice (Figure 2e, f and g), which may explain the neutrophilic infiltration in the brain and meninges of *Nos2*^-/-^ *M*.*tb*-infected mice relative to the C3HeB/FeJ *M*.*tb*-infected mice. As *Nos2*^-/-^ mice displayed a greater severity of CNS-TB disease than C3HeB/FeJ mice in terms of neurobehavior, histopathology, and immunological profile, the *Nos2*^-/-^ mouse strain was chosen for all subsequent experiments.

**Figure 2.**
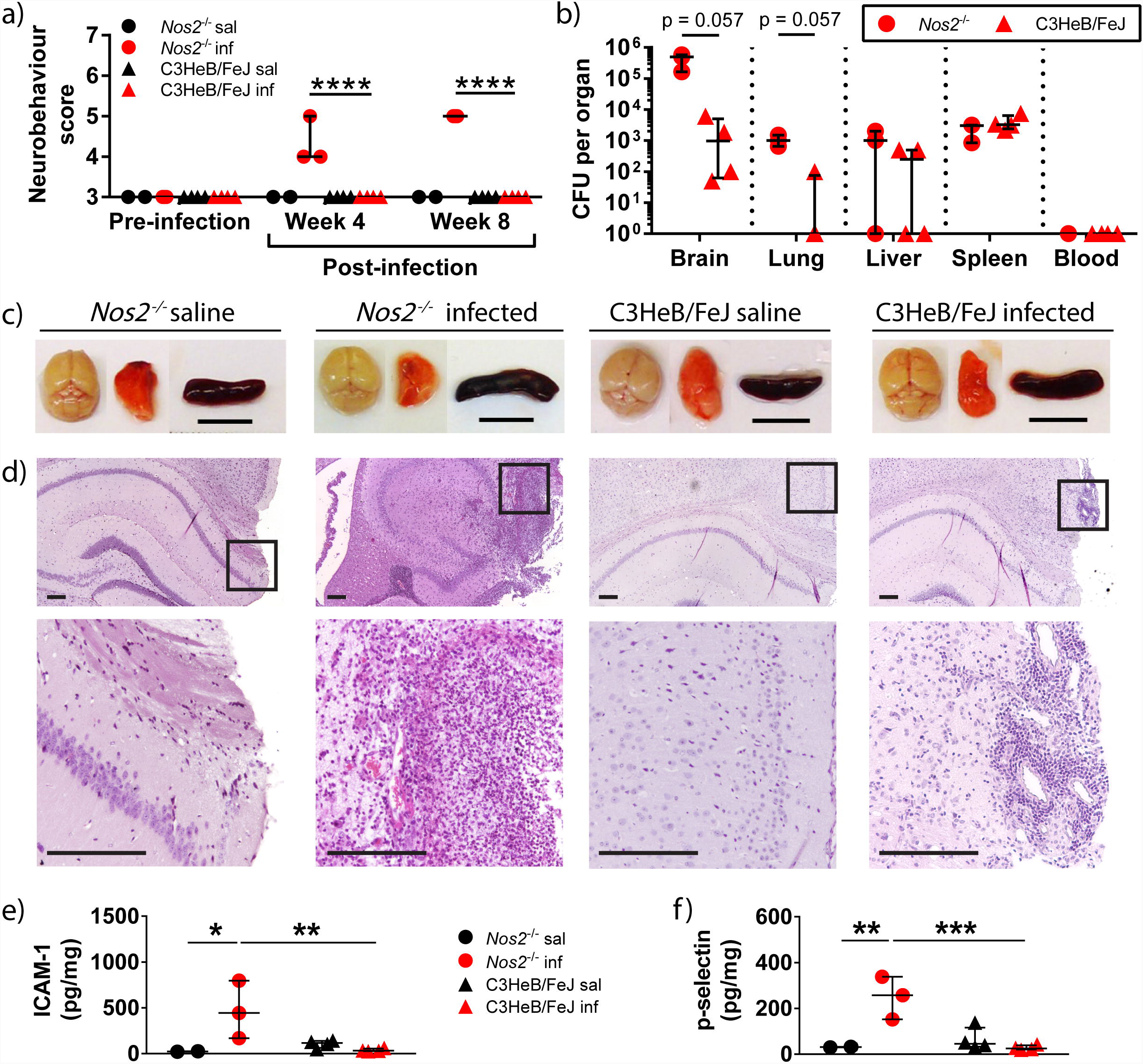
CDC1551 i.c.vent.-infected *Nos2*^-/-^ mice demonstrated increased concentration of Th1-associated cytokines and chemokines, and neutrophil chemoattractants compared to C3HeB/FeJ infected mice at 8 weeks p.i.. Concentrations of chemokines and cytokines in brain homogenates were normalised against total protein concentration. Statistical analysis performed using two way ANOVA with Sidak’s multiple comparisons test. Bars represent median and IQR. *, p < 0.05; **, p < 0.01; ***, p < 0.001. *Nos2*^-/-^ saline: n = 2; *Nos2*^-/-^ infected: n = 3; C3HeB/FeJ saline: n = 4; C3HeB/FeJ infected: n = 4.

### I.c.vent. infection by H37Rv *M*.*tb* strain resulted in a worse neurobehavioral score, earlier mortality and increased mycobacterial load in the brain than CDC1551 *M*.*tb* strain

We further compared two different *M*.*tb* strains, H37Rv and CDC1551, on the neurobehavioral scores and mortality outcomes. At day 28 p.i., infected mice had a significantly lower weight than saline control, independent of the routes of infection (Figure 3a and Supplementary figure 2a). Within the i.c.vent. group, weight change between H37Rv- and CDC1551-infected mice were similar throughout the study (Figure 3a). However, within the i.v. group, the weight change in H37Rv-infected mice at day 28 p.i. was -3.6 ± 3.1% (mean ± s.d.) which was significantly different from CDC1551-infected mice that gained a mean weight of 6.0 ± 2.4% (*p* = 0.0027) (Supplementary figure 2a).

**Figure 3.**
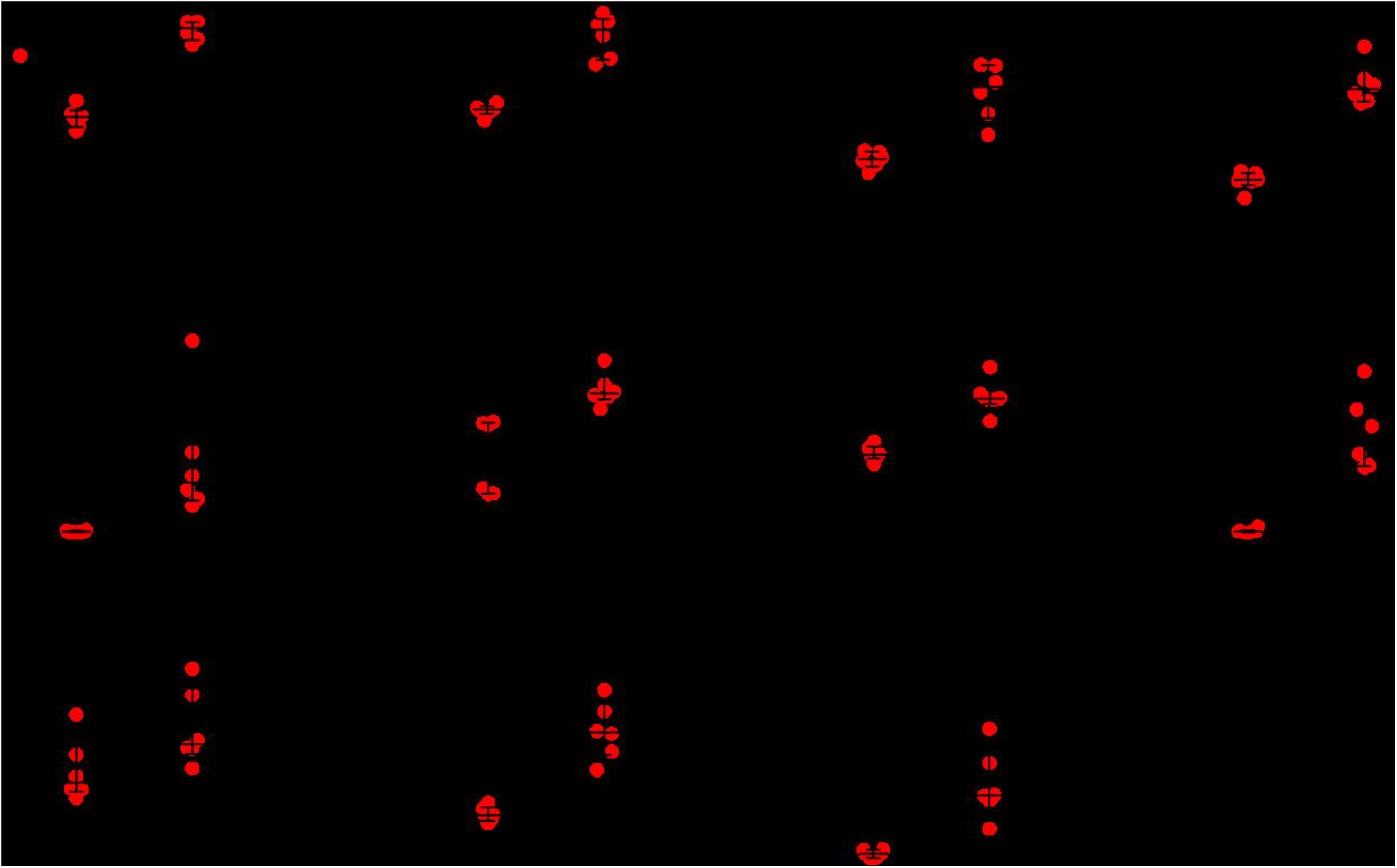
I.c.vent. infection of *Nos2*^-/-^ mice with H37Rv resulted in earlier mortality, higher neurobehavioral score, and increased mycobacterial load in the brain compared to CDC1551. (a) *M*.*tb*-infected mice lost significantly more weight than saline control, analysed using two-way ANOVA with post-hoc Tukey’s multiple comparisons test. Percentage change in body weight relative to initial body weight at day 0 p.i. is shown. Bars represent mean ± SEM. **, p < 0.01; ***, p < 0.001. Statistical analysis between H37Rv-infected mice and saline controls in red asterisks, while comparisons between CDC1551-infected mice and saline controls in blue asterisks. (b) Kaplan-Meier curve shows a significant difference in survival between the groups. (c) H37Rv i.c.vent. demonstrate higher neurobehavioral score at 8 weeks p.i. compared to CDC1551 i.c.vent. mice, analyzed using two-way ANOVA with post-hoc Tukey’s multiple comparisons test. **, p < 0.01; ****, p < 0.0001. (d) Gross pathological examination of brain, lung and spleen show no difference between saline control and *M*.*tb*-infected mice except for enlarged spleen in H37Rv i.c.vent-infected mice. Scale bar = 1 cm. (e) H37Rv-infected mice show a trend towards increased *M*.*tb* load in the brain, while lung and blood CFU were comparable to CDC1551-infected mice. Bars represent median and IQR. Statistical analysis was conducted using Mann-Whitney test. (f) Low-power view of a representative H&E-stained granuloma in the brain of H37Rv i.c.vent. mice. High-power view (inset) demonstrates numerous intra- and extracellular acid-fast bacilli (black arrows) by Ziehl-Neelson (ZN) stain within the brain granuloma. Scale bars represent 1 mm in H&E stain and 20 μm in ZN stain. Saline: n = 6; H37Rv: n = 6; CDC1551: n = 5.

By day 28 p.i., 3 out of 6 (50%) H37Rv i.c.vent.-infected mice were euthanized as they reached the humane end point, compared to 1 out of 5 (20%) in CDC1551-infected mice (Figure 3b). As infection progressed, neurological signs in surviving H37Rv i.c.vent. mice worsened with a higher neurobehavioral score than CDC1551 i.c.vent. mice by week 8 p.i.. Median (IQR) neurobehavioral score in H37Rv i.c.vent. mice was 5.5 (5-6) compared to 4 (4-4) in CDC1551 i.c.vent. infected mice (*p* < 0.0001) (Figure 3c).

Within the group of i.v.-infected mice, H37Rv *M*.*tb* also resulted in higher mortality than CDC1551. *M*.*tb* H37Rv-infected mice displayed uniform lethality by day 30 p.i., while 100% survival was observed in CDC1551-infected mice (Supplementary figure 2b). The findings from the survival curve are also reflected in the neurobehavioral score over time, as CDC1551 i.v. mice displayed mild to no neurological signs at week 8 p.i. (Supplementary figure 2c).

On gross pathology examination, we found that both H37Rv and CDC1551 i.v.-infected mice (Supplementary figure 2d) and H37Rv i.c.vent.-infected mice had enlarged spleen relative to saline controls (Figure 3d), indicating dissemination of infection. CDC1551 i.v.-infected mice developed macroscopic granulomas in the lungs (Supplementary figure 2d) that was not observed in other groups. I.v. inoculation of *M*.*tb*, independent of *M*.*tb* strains, resulted in a disseminated infection with granuloma formation in the heart, kidneys, and spleen, which was not observed in i.c.vent.-infected mice (Supplementary figure 2e). Intra-abdominal abscesses were also found in one of the H37Rv i.v.-infected mice examined (data not shown). This was consistent with the blood culture results, where mycobacteraemia was detected in six out of 12 (50%) mice infected by the i.v. route (Supplementary figure 2f). There was no mycobacteremia in any of the i.c.vent.-infected mice (n = 11) (Figure 3e). H37Rv i.c.vent.-infected mice demonstrated a trend towards increased *M*.*tb* load in the brain than CDC1551 i.c.vent., with median brain CFU count of 4.3 ×10^6^ in H37Rv i.c.vent. and 4.9 ×10^5^ in CDC1551 i.c.vent. mice (*p* = 0.052). Interestingly, although no mycobacteremia was found in i.c.vent.-infected mice, *M*.*tb* was cultured from the lungs with comparable mycobacterial load in both *M*.*tb* strains. Median lung CFU count was 2.9 ×10^4^ and 5.0 ×10^2^ in H37Rv and CDC1551 i.c.vent. mice respectively (*p* = 0.33) (Figure 3e). The presence of *M*.*tb* in the brain was confirmed by Ziehl-Neelsen staining, with numerous intra- and extra-cellular bacilli within the brain granulomatous lesion (Figure 3f).

Collectively, these results showed that H37Rv *M*.*tb* is more suited than CDC1551 *M*.*tb* for the murine CNS-TB model as H37Rv infection resulted in earlier mortality, worse neurobehavioral score and increased mycobacterial load in the brain compared to CDC1551 infection. I.c.vent infection also resulted in a more localized infection relative to the widespread dissemination observed in the i.v-infected mice.

### H37Rv infection via the i.c.vent. route resulted in brain histopathological changes similar to human CNS-TB patients with increased expression of adhesion molecules relative to the i.v. route

We next conducted a thorough histological evaluation in *Nos2*^-/-^ mice infected with H37Rv via either the i.v. or i.c.vent. route. Histopathological evaluation demonstrated that i.c.vent.-infected *Nos2*^-/-^ mice developed more severe meningitis and parenchymal granulomas compared to i.v.-infected mice, independent of *M*.*tb* strains (Figure 4a-c and Supplementary figure 3). More importantly, *M*.*tb*-induced lesions in the brain of H37Rv i.c.vent. mice, which included pyogranuloma, granuloma, and perivascular cuffing (Figure 4d), were similar to the hallmark histological lesions in human CNS-TB patients (Figure 4e). While all infected mice developed brain granulomas independent of the routes of infection and *M*.*tb* strain, pyogranulomas were only present in i.c.vent.-infected mice. These pyogranulomatous lesions contained a central area of liquefactive necrosis with abundant degenerated polymorphs surrounded with MNCs such as macrophages, which were sometimes epithelioid, and lymphocytes enclosed within a thin layer of fibrous capsule (Supplementary figure 4a). These necrotic brain lesions are a key feature in human CNS-TB patients [28]. In addition, the presence of neutrophils in CNS tuberculous granulomas was also demonstrated in human brain biopsies with histologically proven CNS-TB [28]. Other pathological lesions in the brain of H37Rv i.c.vent. mice included gliosis and neuronal degeneration (Supplementary figure 4b). Consistent with the more pronounced brain inflammation, H37Rv i.c.vent.-infected mice had a higher histopathological score than H37Rv i.v. mice (Table 1). In addition, the meningitis and parenchymal inflammation in the brain of H37Rv i.c.vent.-infected mice were extensive, extending far beyond the injection site with a total spread of 2500 μm in the anterior-posterior axis (Supplementary figure 5). Collectively, these results demonstrate that i.c.vent. infection of *Nos2*^-/-^ mice with H37Rv produces a murine CNS-TB model that resembles human necrotic TB granulomas, and also recapitulates the cellular architecture of human CNS-TB tuberculomas.

**Table 1.** Histopathological evaluation of *M*.*tb*-induced lesions in H37Rv-infected *Nos2*^-/-^mice.

| Lesions<sup>A</sup> | H37Rv i.v. |  |  |  | H37Rv i.c.vent. |  |  |  |
| --- | --- | --- | --- | --- | --- | --- | --- | --- |
|  | M | C | H | T | M | C | H | T |
| Inflammation (MNCs) | 0 | 0 | 0 | 0 | 1 | 3 | 2 | 0 |
| Perivascular cuffing |  | 0 | 0 | 0 |  | 3 | 2 | 2 |
| Gliosis |  | 1 | 1 | 0 |  | 3 | 1 | 0 |
| Granuloma |  | 1 | 2 | 0 |  | 2 | 0 | 0 |
| Pyrogranuloma |  | 0 | 0 | 0 |  | 4 | 3 | 0 |
| Neuronal degeneration/necrosis |  | 0 | 0 | 0 |  | 2 | 1 | 0 |
| Liquefactive necrosis (+/-)<sup>B</sup> |  | + | + | - |  | + | + | - |
| Presence of bacilli (+/-)<sup>B</sup> | - | - | + | - | - | + | + | - |

M: meninges; C: cerebral cortex; H: hippocampus; T: thalamus
<sup>A</sup> Severity of lesions in each group are scored on a scale of 0–5: 0 – no abnormalities detected; 1 – minimal; 2 – mild; 3: moderate; 4: marked; 5: severe. The average score of 5–6 mice per group is shown.
<sup>B</sup> +/-: present/absent

**Table 2.** Composite neurobehavioral score criteria for CNS-TB mouse model.

| **Criteria** | **Score** |
| --- | --- |
| **Tremors** |  |
| Absent | 1 |
| Present | 2 |
| **Twitch/jerk** |  |
| Absent | 1 |
| Mild (< 3 in 10 sec) | 2 |
| Severe (≥ 3 in 10 sec) | 3 |
| **Eyes** |  |
| Normal | 1 |
| Closed eyelids | 2 |

**Figure 4.**
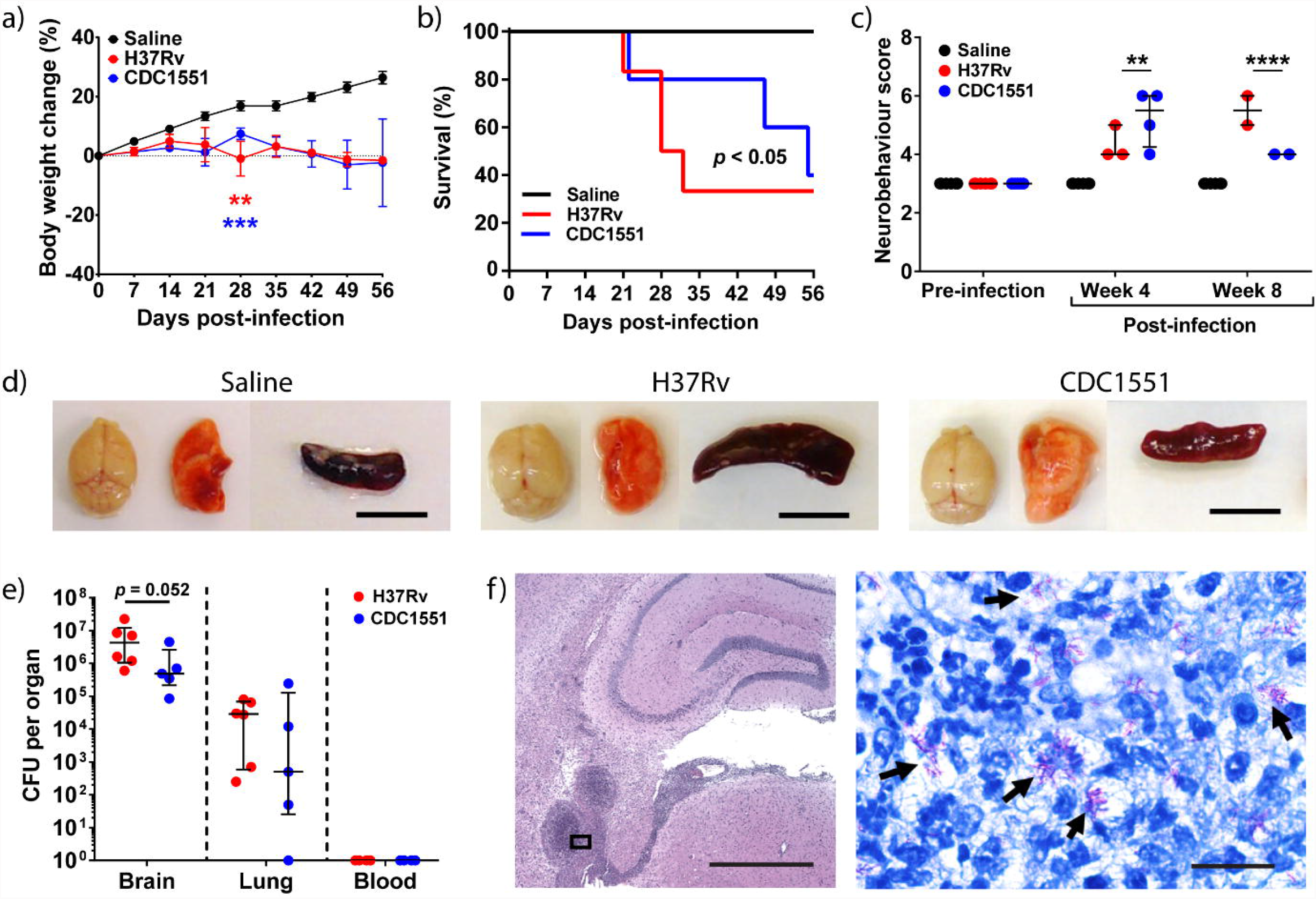
*Nos2*^-/-^ mice infected with H37Rv by the i.c.vent. route developed pyogranulomas and larger granulomatous lesions with increased concentrations of ICAM-1 compared to i.v. route. (a) Overall histopathology via H&E stain demonstrate more severe (b) meningitis and (c) parenchymal granulomatous inflammation in H37Rv i.c.vent. than H37Rv i.v. mice. Bottom panel: high-power views of insets. (d – e) H&E stain demonstrate extensive tissue necrosis (top), granuloma (middle), and perivascular cuffing (bottom) in (d) H37Rv i.c.vent. mice and (e) human CNS-TB biopsy specimen. (f – g) H37Rv i.c.vent.-infected mice had more and larger brain granulomas. The number and size of granulomas in each group were respectively quantified from 6 different sections of 6 mice, analyzed using Mann-Whitney test. H37Rv i.c.vent.-infected mice showed higher levels of (h) ICAM-1 compared to i.v.-infected mice, whereas (i) p-selectin levels were comparable. Bars represent median and IQR. Statistical analysis was conducted using Mann-Whitney test. **, p < 0.01. n = 6 mice were used per experimental group. Scale bar = 1mm (a), 200 μm (b – e).

To analyse the extent of granulomatous inflammation, we measured the number and size of brain granulomas in each group. H37Rv i.c.vent.-infected mice had significantly more granulomas which were larger compared to H37Rv i.v.-infected mice (Figure 4f and g). Median (IQR) granuloma size was 1.18 (0.85-2.18) mm^2^ in H37Rv i.c.vent.-infected mice compared to 0.07 (0.03-0.10) mm^2^ in H37Rv i.v.-infected mice (*p* = 0.0022). Analysis of the adhesion molecules showed that ICAM-1 was significantly increased in i.c.vent.-infected mice relative to i.v.-infected mice (Figure 4g). P-selectin in H37Rv-infected mice was similarly upregulated in both routes of infection compared to saline controls (*p* = 0.0022) (Figure 4h). The higher ICAM-1 expression may explain the increased infiltration of leukocytes which in turn lead to larger granuloma size in the H37Rv i.c.vent.-infected compared to H37Rv i.v.-infected mice.

A similar trend was observed for CDC1551 *M*.*tb* strain. CDC1551 i.c.vent.-infected mice had a higher histopathological score than i.v.-infected mice (Supplementary Table 1), although the number of brain granulomas was similar for both routes of infection with this *M*.*tb* strain (Supplementary figure 6a). The median (IQR) granuloma size in i.c.vent. route of 0.49 (0.43-0.74) mm^2^ was significantly larger than the i.v. route of 0.06 (0.01-0.17) mm^2^ (*p* = 0.0022) (Supplementary figure 6b), with corresponding increase of ICAM-1 expression in the i.c.vent-infected compared to the i.v.-infected mice (Supplementary 6c). In contrast, p-selectin expression was lower in the i.c.vent-infected mice (Supplementary 6d). These findings again indicated that while i.v.-infected mice were capable of developing CNS-TB, the i.c.vent route resulted in a more compartmentalized immunopathological response.

### H37Rv i.c.vent.-infected mice have higher expression of pro-inflammatory cytokines, Th1 chemokines and neutrophil chemoattractants

Inflammatory cytokines found upregulated in the CSF of TBM patients included TNF-α, IFN-γ, IL-1β and IL-6 [29, 30]. To determine if our model has a similar CNS immunological phenotype as human TBM patients, we analysed the expression of pro-inflammatory cytokines in the brain.

Pro-inflammatory cytokines TNF-α, IFN-γ, IL-1β and IL-6 were significantly increased by 17.8-fold, 31.0-fold, 4.8-fold and 7.1-fold, respectively in H37Rv i.c.vent. compared to H37Rv i.v.-infected mice (Figure 5a-d; all *p* < 0.01), and were observed in both *M*.*tb* strains (Supplementary Figure 7a-d). In addition, H37Rv i.c.vent.-infected mice demonstrated 31.7-fold, 7.3-fold, 6.2-fold and 56.8-fold higher expression of Th1 chemokines CCL-3, -4, -5 and CXCL-10 than H37Rv i.v.-infected mice respectively (Figure 5e-h; all *p* < 0.01). This was also observed with the CDC1551 strain (Supplementary Figure 7e-h). The higher concentration of pro-inflammatory cytokines and Th1 chemokines in i.c.vent. mice may explain the more pronounced inflammation and greater extent of inflammatory cell infiltration around cerebral blood vessels in i.c.vent.-infected compared to i.v.-infected mice.

**Figure 5.**
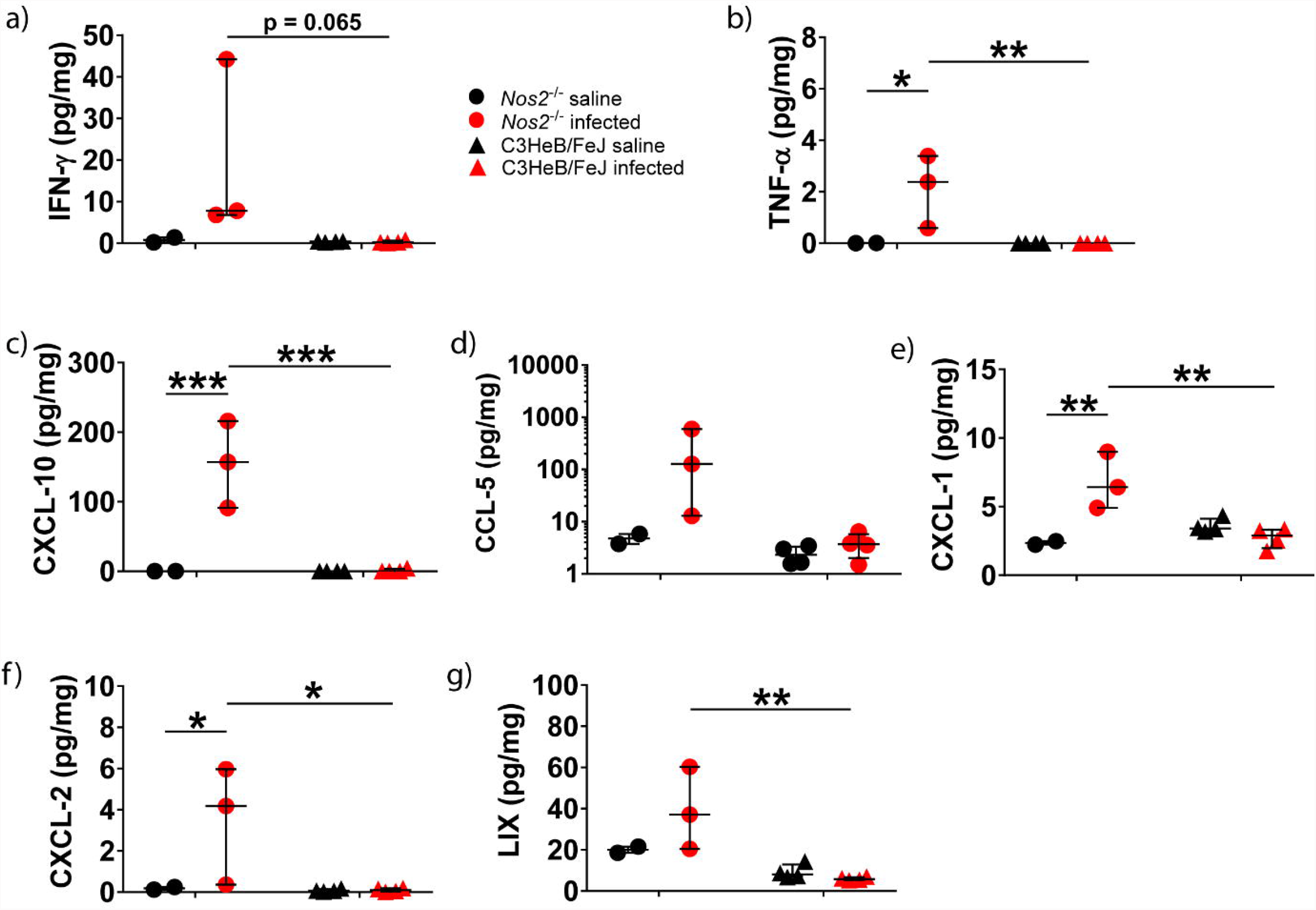
H37Rv infection of *Nos2*^-/-^ mice by the i.c.vent. route resulted in significantly higher brain expression of inflammatory mediators than the i.v. route. H37Rv i.c.vent.- infected mice had higher concentrations of (a-d) pro-inflammatory cytokines, (e-h) Th1 chemokines, and (i-k) neutrophil chemoattractants than H37Rv i.v. mice. Concentrations of inflammatory mediators in the brain were measured after day 21 p.i.. Concentration of each immunological marker was normalised against the total protein concentration. Bars represent median and interquartile ranges. Statistical analysis was conducted using Mann-Whitney test. *, p < 0.05; **, p < 0.01. n = 6 mice were used per experimental group.

Neutrophil chemokines CXCL-1, CXCL-2 and LIX were upregulated by 1.4-fold, 35.6-fold and 5.2-fold, respectively in H37Rv i.c.vent.-infected than H37Rv i.v.-infected mice (Figure 5i-k). While CXCL-1 expression was similar between the two infection routes of CDC1551-infected mice, CDC1551 i.c.vent. mice had higher expression of CXCL-2 and LIX than CDC1551 i.v. mice (Supplementary figure 7i-k). The significantly higher expression of neutrophil chemoattractants in i.c.vent.-infected mice, independent of *M*.*tb* strains, may explain the presence of pyogranulomas with marked neutrophilic infiltration in i.c.vent.- but not i.v.-infected mice.

Collectively, these immunological and histological findings indicate that i.c.vent. infection of *Nos2*^-/-^ mice with H37Rv strain creates a better CNS-TB model than the i.v. route of infection as it exhibited more pronounced brain inflammation as shown by the higher expression of pro-inflammatory cytokines, Th1 chemokines and neutrophil chemoattractants.

## DISCUSSION

Human CNS TB is severe and progress is limited by a lack of good animal model systems that reflect immunopathology in human CNS TB. Our study determined the effects of mouse strain, *M*.*tb* strain and route of infection on the development of a murine CNS-TB model with human-like pathology. Here, we show that i.c.vent. infection of *Nos2*^-/-^ mice with *M*.*tb* H37Rv makes a CNS-TB model that shares similar clinical features of human CNS-TB, including neurological morbidity, high mortality, and increased CNS expression of inflammatory mediators. Importantly, our model demonstrated histological evidence of parenchymal granulomas in the cerebral cortex, hippocampus and the presence of necrotizing granulomas similar to human CNS-TB tuberculomas [31, 32]. The presence of a central area of liquefactive necrosis in pyogranulomas of H37Rv i.c.vent.-infected mice resembled human caseating tuberculomas with central liquefaction, a clinical feature that has not yet been replicated in existing murine CNS-TB models. Other features of human CNS-TB include perivascular infiltration with immune cells and a microglial reaction [31, 33]. Similar to that observed in humans, our CNS-TB model showed the presence of gliosis and perivascular cuffing throughout the brain parenchyma.

We evaluate the simultaneous expression of adhesion molecules, chemokines, and cytokines in an attempt to elucidate the mechanism underlying the chronic inflammatory state in human CNS-TB. While several clinical studies have unanimously demonstrated an increased CSF expression of inflammatory cytokines TNF-α, IFN-γ, IL-1β and IL-6 in TBM patients [17, 29, 30], current murine CNS-TB models have failed to recapitulate this immunological profile [14, 16]. Through immunological analysis, we showed that H37Rv i.c.vent.-infected *Nos2*^-/-^ mice had significantly increased expression of TNF-α, IFN-γ, IL-1β and IL-6, similar to human TBM patients [29, 30], indicating that our pre-clinical model mirrors human CNS-TB. In addition, we demonstrated H37Rv i.c.vent.-infected mice exhibited upregulation of adhesion molecules p-selectin and ICAM-1 compared to saline controls, in keeping with the increased leukocyte infiltration in the brain and extends previous *in vitro* observations [34, 35], and that *M*.*tb* increases expression of endothelial adhesion molecules in a co-culture BBB model [36].

While i.c.vent. infection of *Nos2*^-/-^ mice with either *M*.*tb* H37Rv or CDC1551 resulted in a high mortality (67% and 60% respectively), similar to human CNS-TB [37, 38], H37Rv is superior to CDC1551 as the murine CNS-TB model for two reasons. Firstly, H37Rv infection resulted in the development of more severe neurological deficits with a worse neurobehavioral score and earlier mortality than CDC1551 infection, which reflected the neurological morbidity and severity of disease in human CNS-TB [39, 40]. Secondly, H37Rv-infected mice showed an increased severity of histopathological lesions than CDC1551-infected mice, demonstrated by the greater extent of pyogranulomas and liquefactive necrosis in H37Rv i.c.vent. mice, extending from the cerebral cortex to the hippocampus which were not observed in CDC1551-infected mice, but similar to human CNS-TB histology [28]. This is consistent with previous findings where H37Rv is more virulent than CDC1551 in animal models of pulmonary TB both in rabbits [41] and in mice [42]. Although the mycobacterial load in the brain of H37Rv-infected mice had a trend to increase compared to CDC1551-infected mice, this did not reach statistical significance. A repeat experiment with a lower dose of infection, and a longer experiment with more time points may help to further characterize this CNS-TB model.

Previous murine CNS-TB models have employed direct injection into the brain parenchyma to induce CNS infection [10, 18, 21], which resulted either in granulomas being restricted to the injection site with no widespread inflammation or the absence of granulomas. Thus, to better mimic the rupture of the Rich foci in human CNS-TB, with the subsequent release of *M*.*tb* into the CSF to induce TBM [1], we inoculated *M*.*tb* into the third ventricle for meningeal infection. To prevent surgery-related loss of mice due to excessive bleeding or hemorrhage, we injected *M*.*tb* at an angle into the third ventricle to avoid puncturing the superior sagittal sinus. In addition to the direct CNS inoculation of *M*.*tb* via the i.c.vent. route, we also explored the i.v. route to mimic the natural course of hematogenous spread from the lung to the brain in human CNS-TB [43]. However, we found the i.v. route of infection to be less suited for our murine CNS-TB model, as the mice exhibited a widespread disseminated infection resembling miliary TB, with granulomas observed in multiple organs of the lungs, spleen, heart, and kidneys, but not typical brain lesions. Dissemination of *M*.*tb* to the heart of H37Rv i.v. mice may explain the early and uniform lethality with mortality of these mice by day 30 p.i..

Different mouse strains have different susceptibilities to *M*.*tb* infection, which may explain the varying degree of disease and brain histopathology in existing murine CNS-TB models. To investigate whether the C3HeB/FeJ mice, which are hypersusceptible to pulmonary TB infection [19, 20], or the *Nos2*^-/-^ mice, which have altered innate immune response, are more susceptible to CNS-TB infection, we evaluated the C3HeB/FeJ and *Nos2*^-/-^ mouse strains for our murine CNS-TB model. *M*.*tb*-infected *Nos2*^-/-^ mice exhibited worse neurobehavioral score than C3HeB/FeJ mice and developed neurological symptoms such as myoclonic jerks and limb weakness that resembled seizures and hemiparesis respectively in human CNS-TB patients [1]. In addition, infected *Nos2*^-/-^ mice demonstrated greater inflammatory cell infiltrates, higher expression of adhesion molecules and chemokines in the brain than C3HeB/FeJ mice. Although there was trend to lower mycobacterial load in the C3HeB/FeJ mice, these infected mice expressed similar level of adhesion molecules and chemokines in the brain to saline controls, indicating that the CNS response to infection in the C3HeB/FeJ mice was minimal. These findings show that *Nos2*^-/-^ mice is a better CNS-TB model than C3HeB/FeJ mice, and underscores the role of *Nos2*-induced NO production in inhibiting *M*.*tb* growth in mice [44]. The presence of abundant neutrophilic infiltrates in the brain of *Nos2*^-/-^ mice may be due to the inability of *Nos2*^-/-^ macrophages to contain the *M*.*tb* infection, as NOS2 upregulation by murine macrophages has been shown to be implicated in *M*.*tb* killing [22], which may result in the activation and recruitment of more leukocytes to the site of infection [28]. This may explain the greater extent of brain granulomatous inflammation seen in *Nos2*^-/-^ mice compared to C3HeB/FeJ mice. Nevertheless, the role of NOS2 in *M*.*tb* killing remains controversial as human microglia do not express NOS2 [24, 25] unlike murine microglia [26]. Thus *Nos2*^-/-^ mice may mimic the lack of NOS2 response in human and recapitulate human CNS-TB disease. Future studies investigating *M*.*tb* killing and cytokine and chemokine production by bone marrow-derived macrophages and neutrophils from *Nos2*^-/-^ mice are needed to gain insight into the function and mechanism of NOS2 gene in CNS-TB pathogenesis.

Our study has limitations including the small sample size comparing C3HeB/FeJ and *Nos2*^-/-^ mouse strains in establishing the murine CNS-TB model and the use of only male mice for the study. Nevertheless, our pilot study is useful for formal sample size calculation for future studies. Findings of the CDC1551-infected *Nos2*^-/-^ mice by the i.c.vent route were successfully reproduced in a separate experiment evaluating the *M*.*tb* strain and route of infection for the CNS-TB model. Male mice were used as there is a male predominance in TB [45], and literature has shown males to be more susceptible to mycobacterial infection [46, 47]. Future studies characterising the responses in both genders of mice would be required for application of research findings [48].

## CONCLUSIONS

Altogether, i.c.vent. infection of *Nos2*^-/-^ mice with H37Rv creates a murine CNS-TB model that resembled human CNS-TB immunopathology, exhibiting the worst neurobehavioral score and with a high and early mortality reflecting disease severity and its associated neurological morbidity. In our study, extensive brain inflammation was seen with granulomas and pyogranulomas that resembled the granulomatous inflammation in human CNS-TB patients with a corresponding increase in expression of adhesion molecules, Th1 cytokine response and neutrophil chemoattractants. As this model replicates the histopathological features of human CNS-TB, it is particularly useful for future drug studies to assess the penetration of potential drug candidates into CNS-TB tuberculomas, and evaluate their efficacy in reducing immunopathology and consequently improve neurological outcome in CNS-TB.

## LIST OF ABBREVIATIONS

BCG: *Mycobacterium bovis* BCG
CFU: colony forming units
CNS-TB: Central nervous system tuberculosis
CSF: cerebrospinal fluid
H&E: hematoxylin-eosin
i.c.vent.: intra-cerebroventricular
IFN-γ: Interferon-γ
IL: Interleukin
iNOS: inducible nitric oxide synthase
i.v.: intravenous
*M*.*tb*: *Mycobacterium tuberculosis*
*Nos2*: nitric oxide synthase 2
NO: nitric oxide
p.i.: post-infection
RNI: reactive nitrogen intermediate
*sst1*: super susceptibility to tuberculosis 1
TB: tuberculosis
TBM: Tuberculous meningitis
TNF-α: Tumor necrosis factor-α

## DECLARATIONS

### Ethics approval and consent to participate

The Domain Specific Review Board from National Healthcare Group Singapore approved the study (Reference: 2015/00067) and human brain samples were anonymized for the purpose of this study.

All animal procedures were approved by the Institutional Animal Care and Use Committee of National University of Singapore under protocol R15-1068, in accordance with national guidelines for the care and use of laboratory animals for scientific purposes.

### Consent for publication

Not applicable

### Availability of data and materials

The data sets generated for this study are available on request to the corresponding author.

### Authors’ contributions

C.W.M.O. conceived the study. P.X.Y., H.J.M., M.Q.H. and C.W.M.O. designed the experiments. C.S.L.D. provided and reviewed the human CNS-TB brain biopsy specimens. P.X.Y., H.J.M., M.Q.H., W.Y. and T.P.M. performed the experiments. P.X.Y., R.R. and C.W.M.O. analysed the data. P.X.Y. and C.W.M.O. wrote the first draft of the manuscript which was reviewed and revised by all authors.

## Acknowledgements

The authors would like to thank the operations team of the National University of Singapore BSL-3 core facility for the infrastructure and logistical support of the study. The authors would also like to thank National University of Singapore Comparative Medicine (CM) and the Neuroscience Phenotyping Core (NPC) for animal training and support. The authors are grateful to Professor Paul Elkington and Associate Professor Sylvie Alonso for commenting on the manuscript.

---

**Video 1.** Saline control *Nos2*^-/-^ mice

**Video 2.** *Nos2*^-/-^ mice infected with *M.tb* via the i.c.vent. route exhibited myoclonic jerks and
limb weakness.

