## Supplementary data for "*Nos2*^-/-^ mice infected with *M. tuberculosis* develop neurobehavioral changes and immunopathology mimicking human central nervous system tuberculosis"

**Supplementary table 1. Histopathological evaluation of *M.tb*-induced lesions in CDC1551-infected *Nos2*^-/-^ mice**

| Lesions^A^ | **CDC1551 i.v.** | | | | **CDC1551 i.c.vent.** | | | |
| --- | --- | --- | --- | --- | --- | --- | --- | --- |
|  | M | C | H | T | M | C | H | T |
| Inflammation (MNCs) | 0 | 0 | 0 | 0 | 3 | 3 | 1 | 0 |
| Perivascular cuffing |  | 0 | 0 | 0 |  | 2 | 2 | 1 |
| Gliosis |  | 0 | 0 | 0 |  | 2 | 1 | 0 |
| Granuloma |  | 0 | 1 | 0 |  | 2 | 0 | 0 |
| Pyrogranuloma |  | 0 | 0 | 0 |  | 3 | 0 | 0 |
| Neuronal degeneration/necrosis |  | 0 | 0 | 0 |  | 2 | 1 | 0 |
| Liquefactive necrosis (+/-)^B^ |  | - | - | - |  | + | - | - |
| Presence of bacilli (+/-)^B^ | - | - | + | - | - | + | - | - |

M: meninges; C: cerebral cortex; H: hippocampus; T: thalamus

^A^ Severity of lesions in each group are scored on a scale of 0–5: 0 – no abnormalities detected; 1 – minimal; 2 – mild; 3: moderate; 4: marked; 5: severe. The average score of 5–6 mice per group is shown.

^B^ +/-: present/absent

**Supplementary figure 1. Cannula implantation and experimental timeline.** (a) A schematic representation (coronal section) of the angled cannulation conducted for injection in the third ventricle. Stereotaxic coordinates for injection site from bregma: -1.60 mm posterior, 0 mm lateral, -2.50 mm ventral. Coordinates for drilling site on skull from bregma: -1.60 mm posterior, 0.80 mm lateral (left). Guide cannula is inserted at an angle of 17.7ᴼ. (b) A schematic representation (sagittal section) of the ventricular system in the mouse brain showing the approximate positions of the lateral ventricle (LV), third-ventricle (3V), fourth-ventricle (4V) and aqueduct (Aq). The 4 dotted lines indicate the approximate location of the brain images shown in (c) with the stereotaxic coordinates posterior from Bregma annotated below. (c) Images of coronal sections of the brain after trypan blue dye administration. If cannula is successfully implanted in the third ventricle, the trypan blue dye will be distributed throughout the ventricular system as demonstrated in the four brain coronal sections. Images shown are representative of four cannulated mice. (d) A schematic representation of the experimental timeline for mice infected with *Mycobacterium tuberculosis* (*M.tb*) via the intra-cerebroventricular route. (e) Low-power view of a representative H&E-stained inflammatory lesion in the brain of C3HeB/FeJ mice infected with 10^5^ CFU of *M.tb* CDC1551. High-power view (inset) demonstrates marked meningeal inflammation indicating a successful ventricular infection with sparing of the brain parenchyma. Scale bars represent 500 µm in low-power view and 100 µm in high-power view. Image representative of n = 4 C3HeB/FeJ mice

**Supplementary figure 2. *Nos2*^-/-^ mice infected with *M.tb* via the i.v. route demonstrate disseminated granulomas with mycobacteraemia in 50% of mice.** (a) *M.tb*-infected mice lost significantly more weight than saline control. At day 28 p.i., H37Rv i.v. mice lost more weight than CDC1551 i.v. mice (§§). Percentage change in body weight relative to initial body weight at day 0 p.i. is shown. Bars represent mean ± SEM. *, p < 0.05; §§, p < 0.01; ****, p < 0.0001. Statistical analysis between H37Rv-infected mice and saline controls in red asterisks, while comparisons between CDC1551-infected mice and saline controls in blue asterisks. (b) Kaplan-Meier curve shows a significant difference in survival between the groups. (c) I.v.-infected mice demonstrate similar neurobehavioral score at 4 weeks p.i.. No data is available for H37Rv i.v. at 8 weeks p.i. as all mice died or reached humane endpoints by day 30 p.i.. (d) CDC1551 i.v.-infected mice developed granulomas (circled) in the lungs. Gross pathological examination of the brain, lung and spleen 21 days after infection. Images are representative of 5-6 mice per condition. (e) Granulomas (circled) are present in the kidneys, heart and spleen of i.v.-infected mice. Scale bars represent 1 cm. (f) H37Rv-infected mice demonstrate increased mycobacteraemia, while brain and lung CFU were comparable to CDC1551-infected mice. Bars represent median and IQR.

**Supplementary figure 3. *Nos2*^-/-^ mice infected with CDC1551 by the i.c.vent. route had more severe meningitis and granulomas compared to i.v. route.** (a) Overall histopathology, (b) meningeal inflammation and (c) parenchymal granulomas are shown. Bottom panel: high-power views of insets. Histology is representative of 5-6 mice. (a) Scale bar = 1 mm. (b) and (c) Scale bar = 200 µm.

**Supplementary figure 4. Pathological lesions in the brain of H37Rv i.c.vent.-infected *Nos2^-/-^* mice.** (a) Well-formed pyogranuloma (P) in the hippocampus surrounded with sheets of inflammatory infiltrate (line) and covered with thin fibrous capsule (arrow). High-power view (inset) of the pyogranuloma shows presence of degenerating neutrophils (DN) at the centre, surrounded with macrophages (M), epithelioid cells (E) and few lymphocytes (L). Scale bar = 200 µm (20 µm in high-power view). (b) Gliosis (black arrow) and neuronal necrosis present with pyknotic nucleus and eosinophilic cytoplasm (blue arrow). Scale bar = 50 µm. Histology representative of 6 mice.

**Supplementary figure 5. Extensive meningeal and parenchymal inflammation observed in the brain of *Nos2*^-/-^ mice infected with H37Rv by the i.c.vent. route.** Serial H&E-stained histopathological sections (each 500 µm apart) of the brain from the same animal shows the granuloma development in the anterior-posterior axis. Scale bar = 1 mm. Histology representative of 5-6 mice.

**Supplementary figure 6. CDC1551 i.c.vent.-infected *Nos2*^-/-^ mice had larger granulomas with increased concentration of adhesion molecule ICAM-1 than CDC1551 i.v. mice.** (a) CDC1551-infected mice by the two routes of infection demonstrate similar number of brain granulomas. (b) CDC1551 i.c.vent.-infected mice had larger granulomas than CDC1551 i.v.-infected mice. The number and size of granulomas in each group were respectively quantified from 6 different sections of 5-6 mice. CDC1551 i.c.vent.-infected mice show higher expression of (c) ICAM-1 compared to i.v.-infected mice, whereas (d) p-selectin expression was higher in i.v.-infected than i.c.vent.-infected mice. Bars represent median and IQR. Statistical analysis was conducted using Mann-Whitney test. **, p < 0.01.

**Supplementary figure 7. CDC1551 infection of *Nos2*^-/-^ mice by the i.c.vent. route resulted in significantly higher brain expression of inflammatory mediators than i.v. route.** CDC1551 i.c.vent.-infected mice had higher expression of (a-d) pro-inflammatory cytokines and (e-h) Th1 chemokines than CDC1551 i.v. mice. Among neutrophil chemoattractants, (i) CXCL-1 expression was similar between the two infection routes, while (j) CXCL-2 and (k) LIX were significantly increased in CDC1551 i.c.vent. than CDC1551 i.v. mice. Inflammatory mediators in the brain were measured after day 21 p.i.. Concentration of each immunological marker was normalised against the total protein concentration. Bars represent median and interquartile ranges. Statistical analysis was conducted using Mann-Whitney test. **, p < 0.01.
