## Supplementary figures and images for "*Nos2*^-/-^ mice infected with *M. tuberculosis* develop neurobehavioral changes and immunopathology mimicking human central nervous system tuberculosis"

### Supplementary figure 1

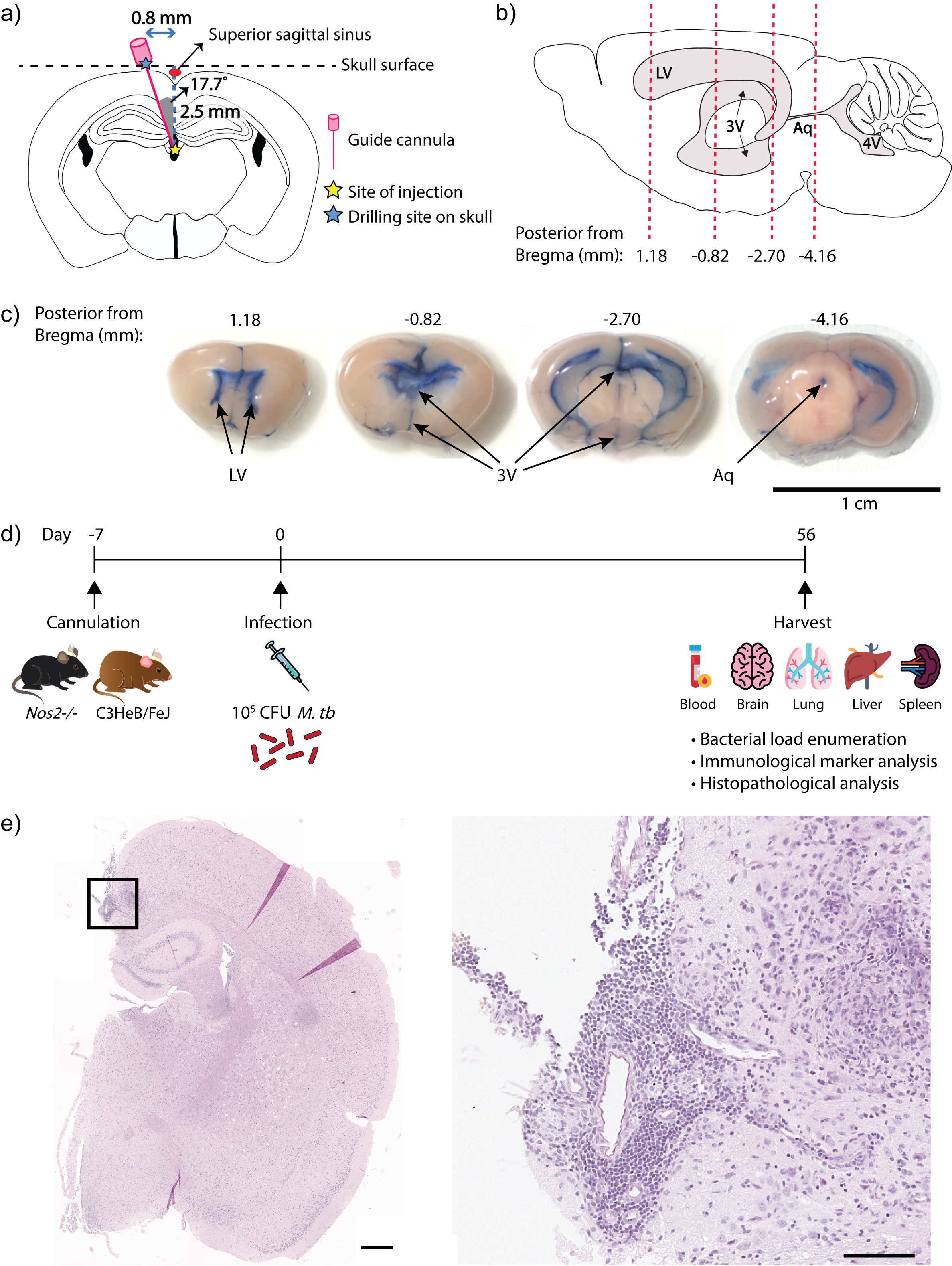

### Supplementary figure 2

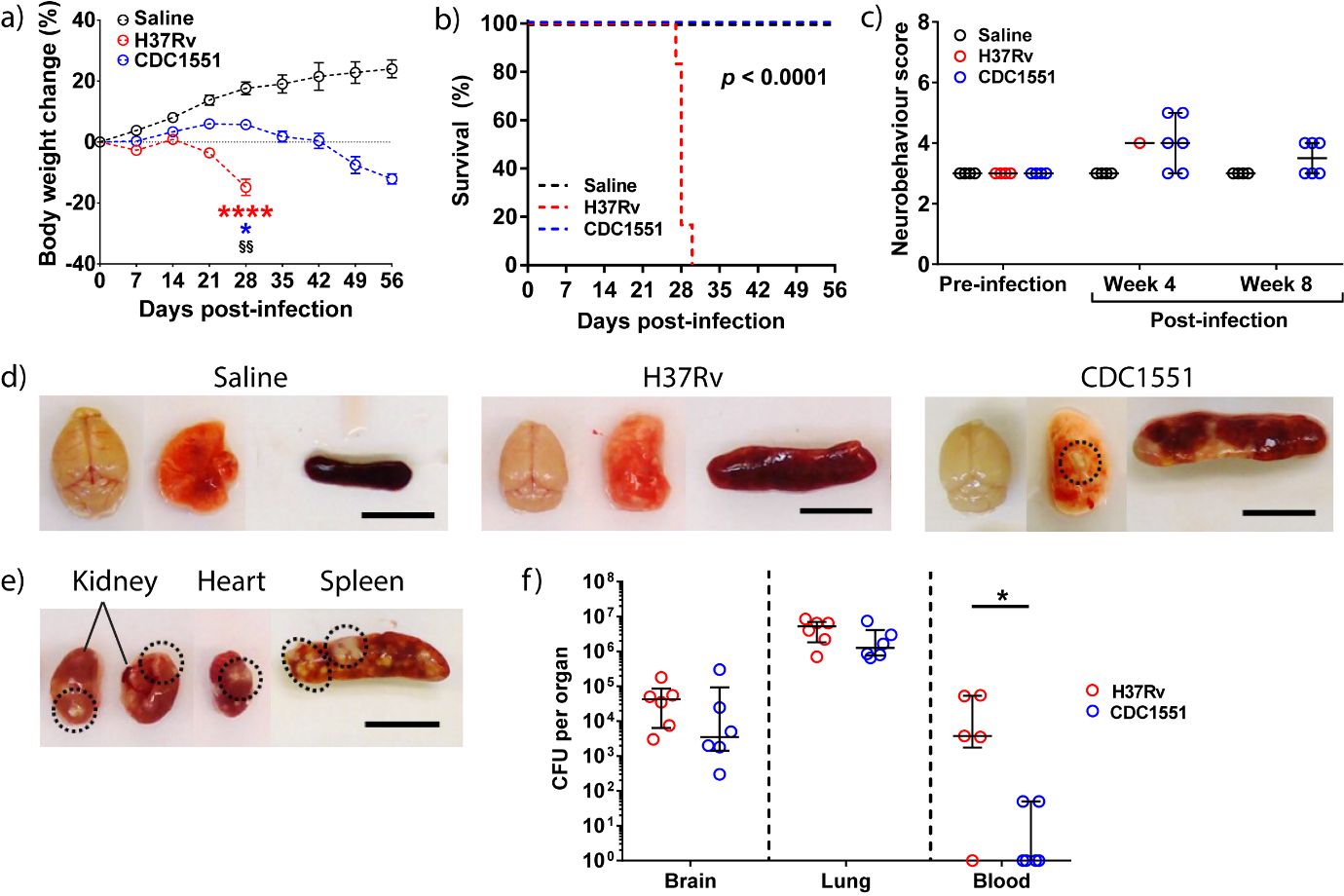

### Supplementary figure 3

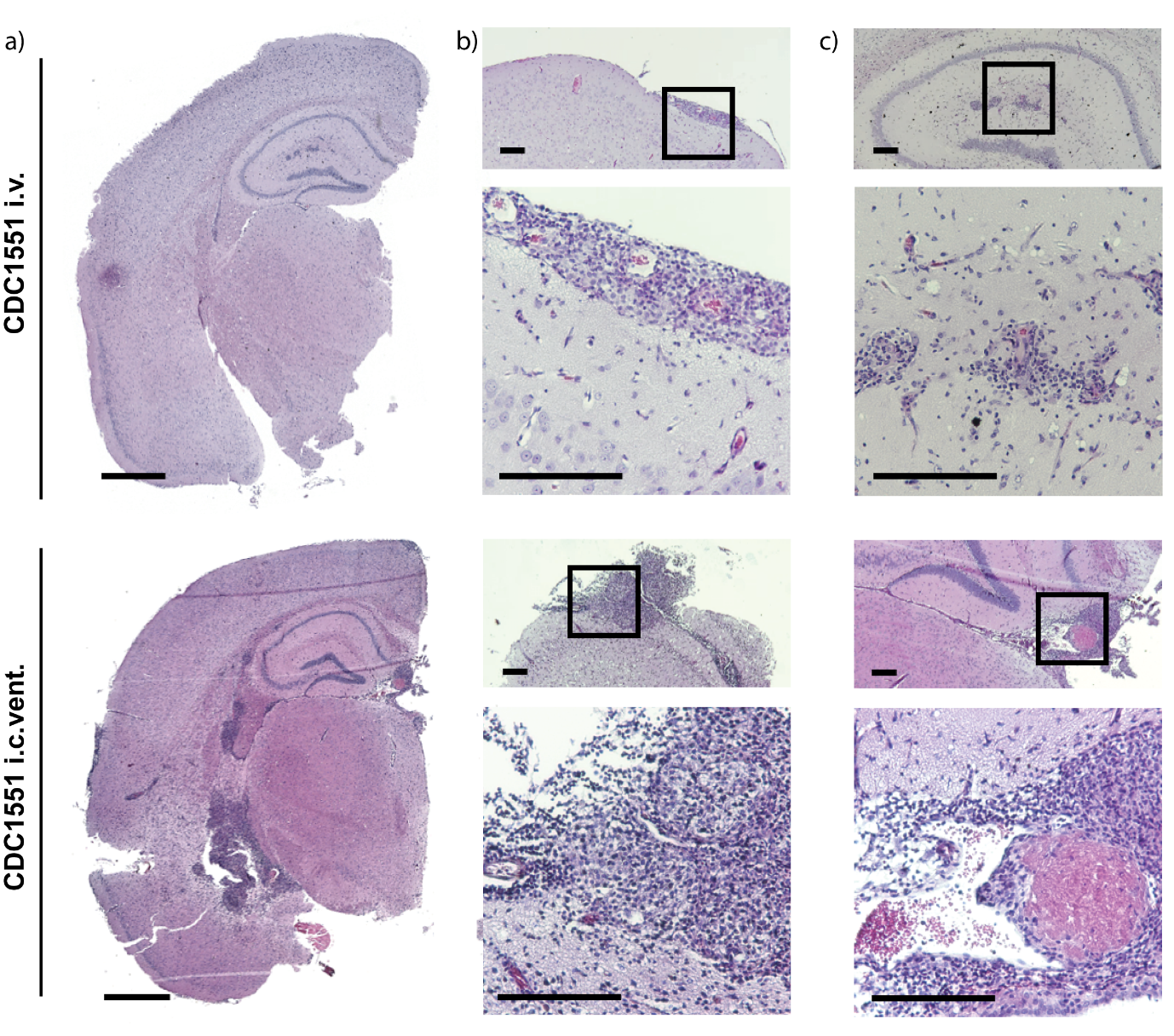

### Supplementary figure 4

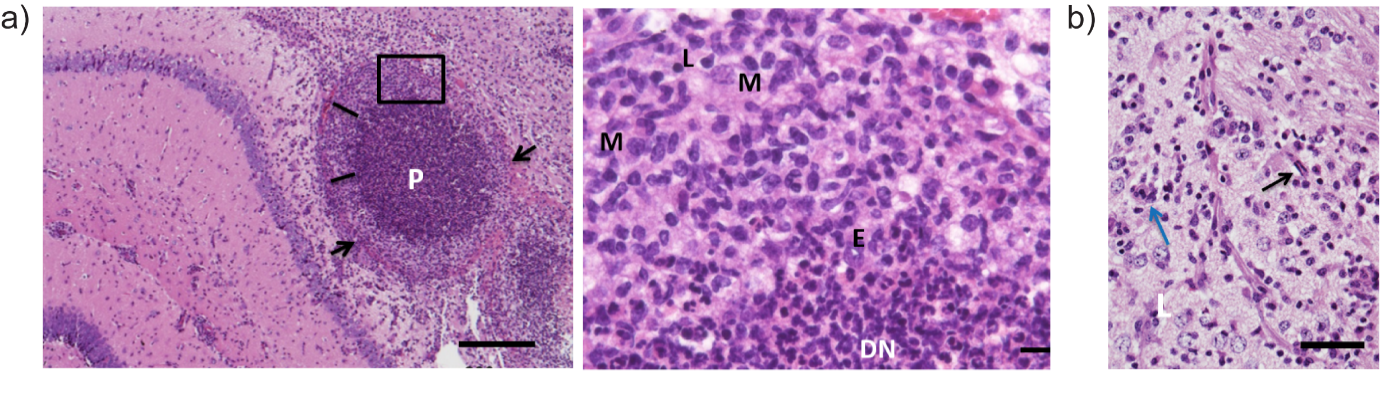

### Supplementary figure 5

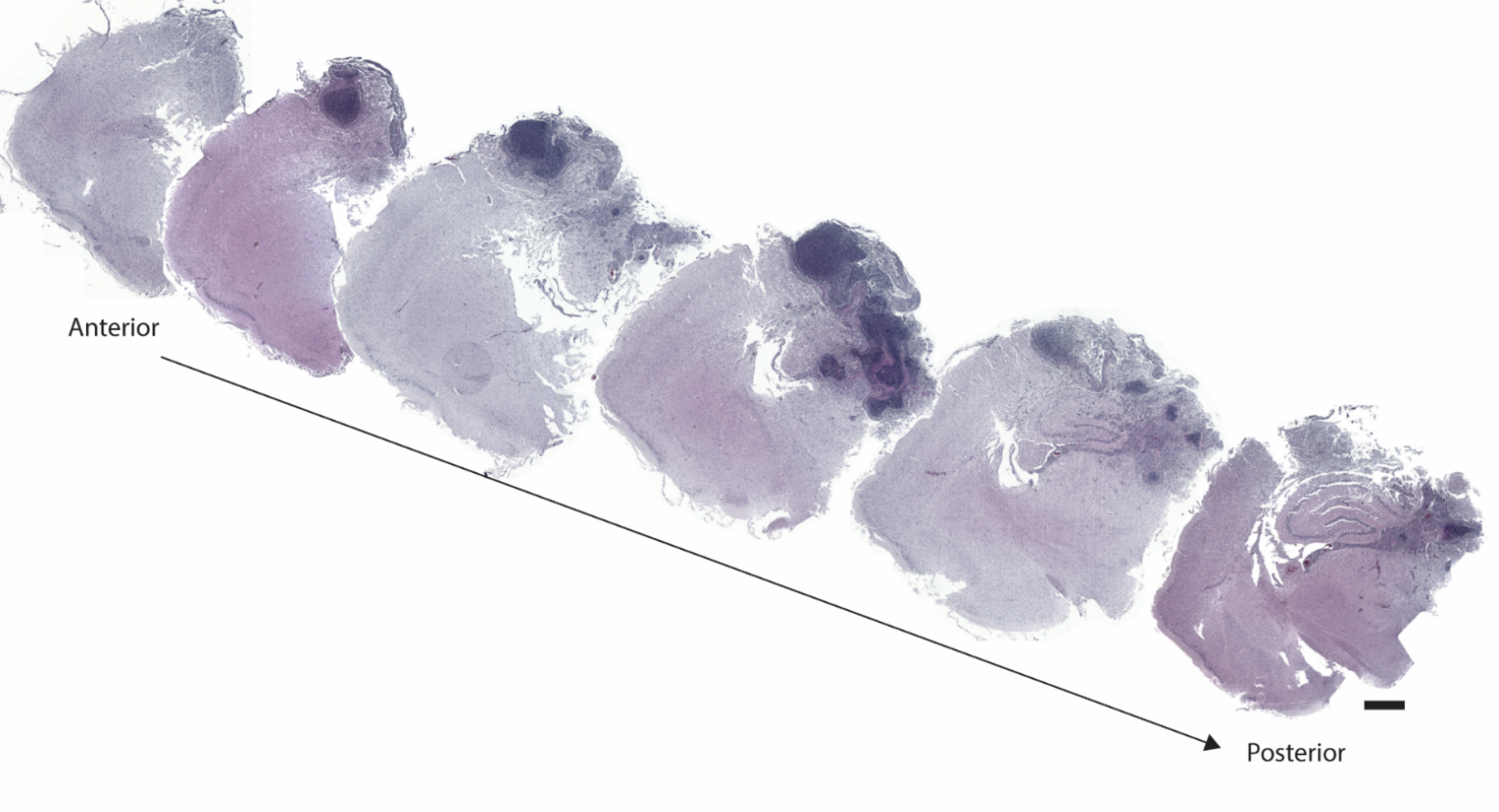

### Supplementary figure 6

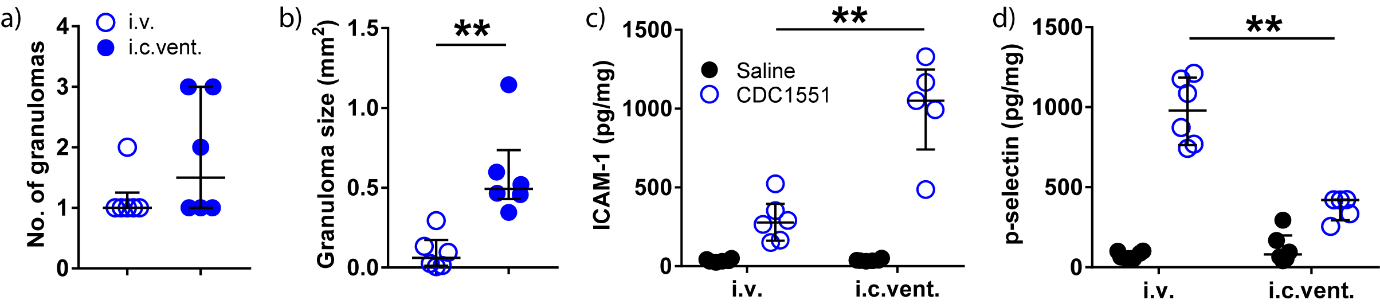

### Supplementary figure 7

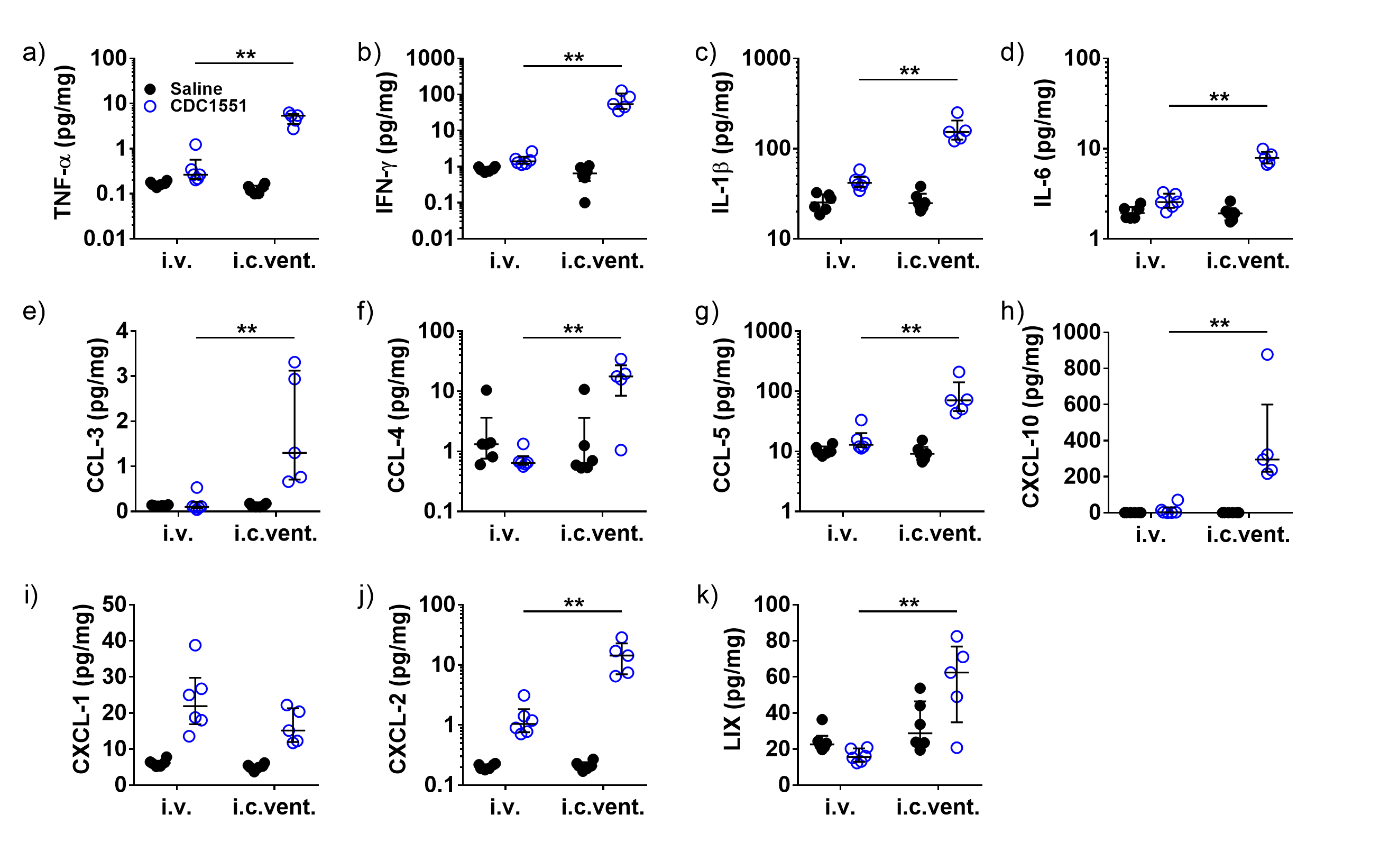
